# Cancer-type specific aneuploidies hard-wire chromosome-wide gene expression patterns of their tissue of origin

**DOI:** 10.1101/563858

**Authors:** Noam Auslander, Kerstin Heselmeyer-Haddad, Sushant Patkar, Daniela Hirsch, Jordi Camps, Markus Brown, Daniel Bronder, Wei-Dong Chen, Rachel Lokanga, Darawalee Wangsa, Danny Wangsa, Yue Hu, Annette Lischka, Rüdiger Braun, Georg Emons, B. Michael Ghadimi, Jochen Gaedcke, Marian Grade, Cristina Montagna, Yuri Lazebnik, Michael J. Difilippantonio, Jens K. Habermann, Gert Auer, Eytan Ruppin, Thomas Ried

**Author notes:** equal contribution first authors (Noam Auslander, Kerstin Heselmeyer-Haddad, Sushant Patkar). equal contribution last authors (Eytan Ruppin, Thomas Ried).

## Abstract

Most carcinomas have characteristic chromosomal aneuploidies specific to the tissue of tumor origin. The reason for this specificity is unknown. As aneuploidies directly affect gene expression, we hypothesized that cancer-type specific aneuploidies, which emerge at early stages of tumor evolution, confer adaptive advantages to the physiological requirements of the tissue of origin. To test this hypothesis, we compared chromosomal aneuploidies reported in the TCGA database to chromosome arm-wide gene expression levels of normal tissues from the GTEx database. We find that cancer-type specific chromosomal aneuploidies mirror differential gene expression levels specific to the respective normal tissues which cannot be explained by copy number alterations of resident cancer driver genes. We show that cancer-type specific aneuploidies “hard-wire” chromosome arm-wide gene expression levels present in normal tissues and propose that the clonal evolution of cancer is initiated by tissue-specific transcriptional requirements.

---

The generally accepted concept of tumorigenesis is based on the notion that genetic alterations that result in malignant transformation are *a priori* tumor promoting. These alterations occur as activating mutations or amplifications of proto-oncogenes, inactivating mutations or deletions of tumor suppressor genes, and chromosomal translocations, that result in the constitutive activation of proliferation-promoting or the inhibition of anti-cell death pathways^1^. The initial driver of tumorigenesis would therefore be “oncogenic” and inherent to the emerging cancer cell.

In solid tumors of epithelial origin, i.e., carcinomas, and in certain other solid tumors such as glioblastoma multiforme and malignant melanoma, aneuploidies of specific chromosomes define the landscape of somatically acquired genetic changes^2–6^. Remarkably, the distribution of ensuing genomic imbalances is strictly cancer-type specific^6,7^. For instance, colorectal carcinomas are defined by extra copies of chromosomes and chromosome arms 7, 8q, 13q and 20q, accompanied by losses of 8p, 17p and 18q^8^. In contrast, cervical carcinomas invariably carry gains of chromosome arms 1q and 3q. In other words, a gain of 3q is not observed in colorectal cancer, and cervical carcinomas do not have copy number gains of, e.g., chromosomes 7 or 13q (Fig. 1A). Tissue-specific chromosomal aneuploidies emerge in dysplastic, i.e., not yet malignant, lesions (Fig. 1B), that, when these aneuploidies are present, are prone to progress to invasive disease^9,10^.

**Figure 1:**
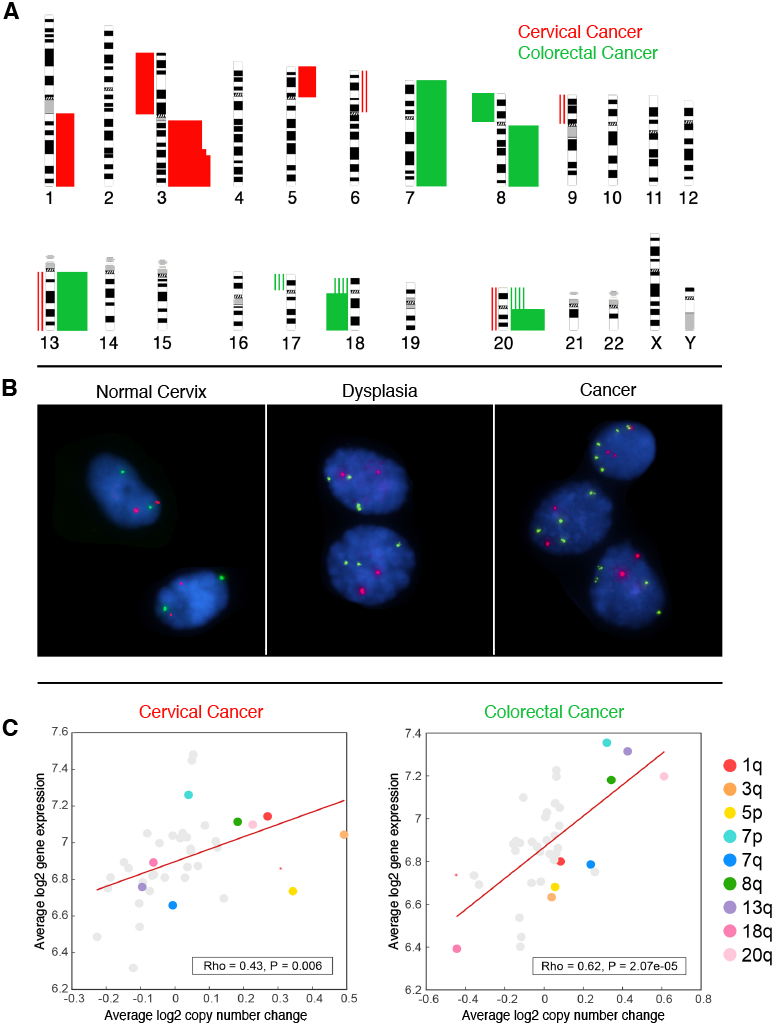
Chromosomal aneuploidies and genomic imbalances and the consequences on the transcriptome of cancer cells. A: Genomic imbalances specific for cervical (red) and colorectal (green) carcinomas. Gains of chromosomes or chromosome arms are presented as bars on the right side of the ideograms, losses on the left. Changes that are visualized as solid blocks occurred in more than 50% of analyzed cases. The distribution of imbalances allows discernment of the tumor entities. B: Interphase FISH analysis of the sequential clonal copy number gain of chromosome arm 3q during cervical tumorigenesis. The FISH analysis shows diploid copy numbers for the probes targeting the centromere of chromosome 7 (red), and the human telomerase gene *TERC* on chromosome arm 3q (green) in normal cervical cells. The gain of chromosome arm 3q results in the clonal expansion of cells in dysplastic lesions (3 copies in all cells), and continues to be gained or even amplified in invasive carcinomas despite increased chromosomal instability and intratumor heterogeneity. C: Genomic copy number changes affect resident gene expression levels. Transcripts from genes on chromosomes that are recurrently gained are more abundant, those on lost chromosomes less abundant. The analysis reflects the TCGA dataset.

The cancer-type specific distribution of genomic imbalances was recently confirmed in two comprehensive pan-cancer analyses of several thousand tumors^11,12^. The pattern of chromosome-arm and whole chromosome gains and losses allows classification of tumor entities. Of note, this distinctive power dissipates when solely considering focal copy number alterations (defined as copy number alterations with lengths < 0.5 chromosome arms) including, but not limited to, presumed or known oncogenes or tumor suppressor genes^5,11^.

The reason for the remarkable cancer-type and tissue specificity of chromosomal aneuploidies is not known. What is well known, however, is that chromosome-wide alterations of gene expression levels follow genomic copy number changes^13,14^, i.e., the transcripts of genes that are located on gained chromosomes are more, and those on lost chromosomes are less abundant. This correlation has been firmly established in primary human carcinomas, in derived cell lines, and in experimental models^13,15–18^. Given the direct effect of genomic copy number on gene expression, chromosomal aneuploidies are therefore a mechanism by which gene dosage can be altered in a “hard-wired” fashion to persist in subsequent generations. This correlation is exemplarily shown for colorectal and cervical cancer based on TCGA data (Fig. 1C)^19,20^.

We therefore hypothesized that cancer-type specific chromosomal aneuploidies create a transcriptional landscape beneficial for cells in the respective normal tissue of origin. Our hypothesis predicts that cancer-specific chromosomal imbalances mirror gene expression patterns in the normal tissues of origin. For example, we surmise that the gain of chromosomes 7 and 13q in colorectal cancers means that genes located on these chromosomes are expressed in normal colorectal tissue more abundantly than in other normal tissues. Accordingly, tissue-specific aneuploidies at early stages of tumorigenesis may not reflect an *a priori* oncogenic stimulus, but rather manifest a proliferative advantage triggered by the physiological requirements of the respective normal tissue.

Assuming random chromosome segregation errors in different tissues, cells that gain chromosomes required for physiological function would be selected for in the context of the respective tissues, triggering a “benign” clonal expansion. The increasing pool of such cells, associated with higher tissue-specific fitness, higher proliferative activity or less cell-death, however, may increase the risk for subsequent genetic damage^21^, while maintaining the overexpression of genes that are required for the physiology of the organ.

To verify or falsify our hypothesis, we analyzed whether cancer-type specific patterns of chromosomal aneuploidies correlate with patterns of chromosome arm-wide gene expression in the respective normal tissues. If the hypothesis is valid, one would expect, for instance, that genes on chromosome arm 13q, which is frequently gained in colorectal, but not in other cancers, are expressed higher in *normal* colorectal epithelium compared to other *normal* tissues.

## RESULTS

### Comparison of cancer-type specific chromosome arm-wide genomic imbalances with the chromosome arm-wide gene expression levels in their respective normal tissues

Copy number alterations based on the TCGA database were extracted from Taylor and colleagues^12^ for 15 different tumor entities and are consistent with previous results^2,3,6,11^. Gene expression levels in the respective normal tissues were retrieved from the GTEx database^22^. We found that, consistent with our hypothesis, across distinct cancer entities, gene expression levels in normal tissues are upregulated on those chromosomes that were gained in the respective tumors, whereas expression levels were lower on lost chromosomes (Fig. 2A). The correlations of arm-level copy number changes in the tumors and the arm-wide gene expression levels in the corresponding normal tissues were invariably positive (Fig. 2B,C). With the exception of acute myeloid leukemia, a tumor without recurrent copy number changes (see Fig. 2A), the correlations were statistically highly significant, and could not be obtained with randomly shuffled data (empirical P-value <0.001).

**Figure 2:**
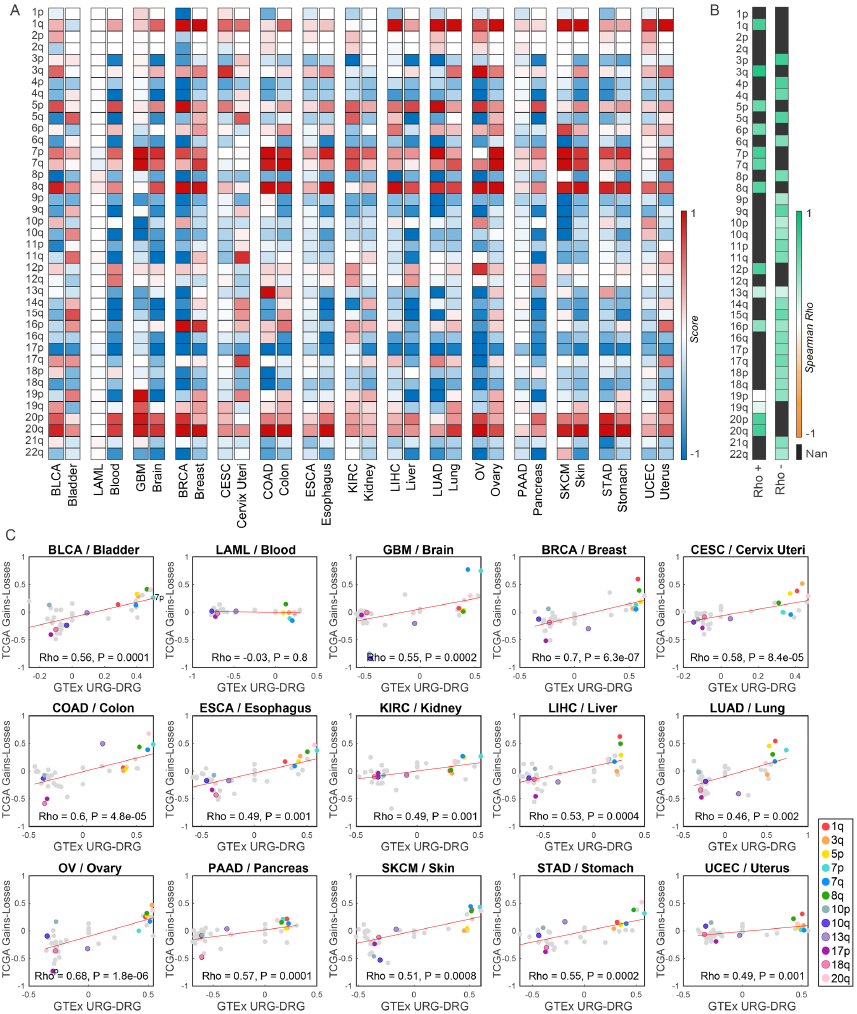
A: Correlation of chromosome arm-wide copy number levels in 15 tumor entities (left column) and arm-wide gene expression levels in the respective normal tissues (right column), respectively. Red, gains; blue, losses. The hue of the colors indicates the frequency of copy number changes and the level of chromosome-arm wide gene expression changes, respectively. BLCA, bladder cancer; LAML, acute myelocytic leukemia; GBM, glioblastoma multiforme; BRCA, breast cancer; CESC, cervical squamous carcinoma; COAD, colorectal adenocarcinoma; ESCA, esophageal squamous carcinoma; KIRC, renal cell carcinoma; LIHC, hepatocellular carcinoma; OV, ovary carcinoma; PAAD, pancreatic adenocarcinoma; SKCM, melanoma; STAD, stomach adenocarcinoma; UCEC, uterine endometrial carcinoma. B: Significance of correlation between the frequency of the gain of each arm in every cancer type, and the number of up-regulated genes in the corresponding normal tissue (Rho+), and the frequency of the loss of each arm in every cancer type, and the number of down-regulated genes in the corresponding normal tissue (Rho-). Black (Nan) indicates those chromosomes that were neither gained (rho+) or lost (rho-) in any of the tumors (with the threshold defined in Materials and Methods) and therefore not included in the analysis. C: Correlation coefficient and statistical significance between chromosome-arm wide copy number changes (Y-axis) and arm-wide gene expression in respective normal tissues (x-axis). With the exception of LAML (no copy number changes), all correlations are highly significant. Relevant chromosome arms are indicated with different colors.

### Classification of normal tissues by chromosome arm-wide gene expression levels

Our hypothesis predicts that if cancers can be identified by cancer-type specific aneuploidies, then normal tissues can be identified by the patterns of gene expression from the corresponding chromosome arms. Hence, we asked whether chromosome arm-wide gene expression levels can classify normal tissues. To this end, we used the median expression levels of each chromosome arm, and applied K-Nearest-Neighbors (KNN) multi-tissue classification with leave-one-out cross validation and principal component analysis. We found that arm-level gene expression in normal tissues could classify tissue-type with high accuracy for almost all tissues (exceptions occurred when the sample size was too small for KNN)(Fig. 3A, Supplementary Fig. 1). These results could not be obtained when randomly assigning genes to chromosomes (empirical P-value < 0.001). Moreover, chromosome arm-wide median gene expression levels were a better predictor of normal tissue origin than global gene expression profiles.

**Figure 3:**
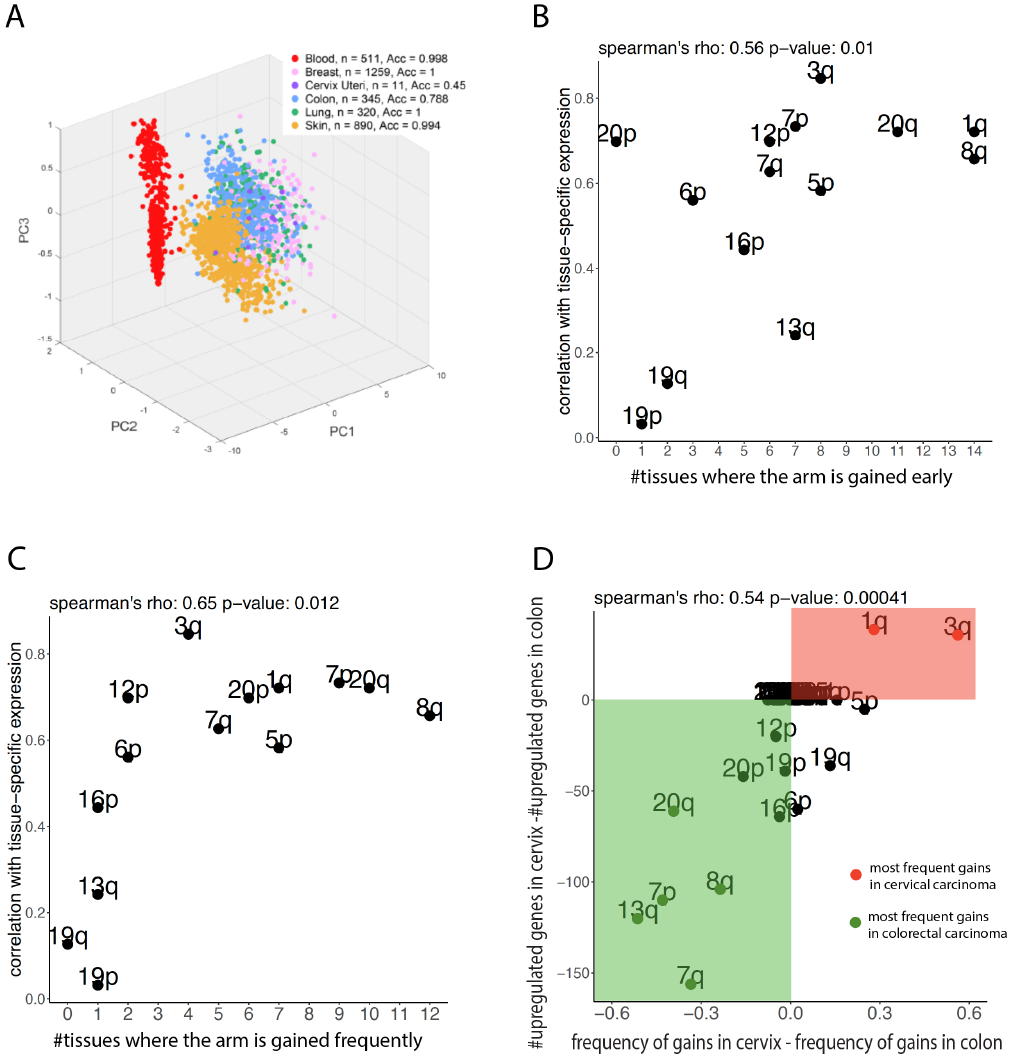
A: Chromosome arm-wide gene expression profiles allow tissue classification. The PCA analysis of chromosome arm-wide gene expression of normal tissues is displayed for six tissues analyzed for clarity. The different normal tissues can be predicted with high accuracy. B: Chromosome arms that are gained early in tumorigenesis show a stronger correlation with chromosome arm-wide gene expression levels in normal tissues. X-axis: number of tissues for which a chromosome arm is gained early (Materials and Methods), Y axis: correlation of arm gain with tissue specific expression (Rho+; Figure 2B) C: Chromosome arms that are gained more frequently across tumor entities show a stronger correlation with chromosome arm-wide gene expression levels in normal tissues. An arm was defined to be frequently gained in a tissue if it was among the top 5 most frequently gained arms in that tissue. X-axis: number of tissues for which a chromosome arm is frequently gained, Y axis: correlation of arm gain with tissue specific expression (Rho+; Figure 2B) D: Comparison of chromosome arm aneuploidies in cervical colorectal cancer with respective gene expression levels in normal cervical and colorectal epithelium. More genes on chromosome arms 1q and 3q are upregulated in normal cervix (red) than in normal colorectum while more genes on, i.e., 7, 8q, 13q and 20q are upregulated in normal colorectum compared to normal cervix.

After having established that (i) cancer-type specific chromosomal aneuploidies mirror chromosome arm-wide gene expression levels in the respective normal tissues and (ii) that these expression levels can predict the tissue of origin, we asked whether those aneuploidies that occur earlier in tumorigenesis have a stronger correlation with normal tissue specific gene expression compared to late events. Hence, the timing of changes in each tissue was computationally determined (Materials and Methods). Indeed, across the tumors analyzed, earlier changes matched tissue-specific chromosome arm-wide gene expression levels better than later events (Fig. 3B). We also showed that frequently gained chromosomes across cancer types correlate with chromosome arm-wide gene expression level in the respective normal tissues better than chromosomes that are rarely gained (Fig. 3C). Finally, we compared chromosome arm-wide gains in cervical and colorectal cancer with chromosome arm-wide gene expression levels in the respective normal tissues. The results confirm our findings: more genes on chromosome arms 1q and 3q, which are gained at early stages of cervical tumorigenesis, are expressed in normal cervix compared to normal colon (Fig. 3D).

### Cancer-type specific aneuploidies cannot be explained by the chromosome arm-wide distribution of cancer genes and methylation patterns

One possible explanation for the correlation of cancer-type specific chromosomal aneuploidies with chromosome arm-wide gene expression levels in the respective normal tissues could be the distribution of oncogenes and tumor suppressor genes^5^. We therefore asked whether the chromosome-arm wide distribution of oncogenes and tumor suppressor genes known to be involved in different tumor types correlates with the acquisition of cancer-type specific imbalances. For the 15 tumor types analyzed we found that the gain or loss of oncogenes or tumor suppressor genes, respectively, does not match the patterns of chromosome arm-wide copy number changes observed in the corresponding tumors (Supplementary Fig. 2). This implies that the distribution of cancer-type genomic imbalances is likely not a reflection of specific cancer genes acting in the respective tumors, which is in accordance with previous results obtained by Beroukhim and colleagues^11^. Potential mechanisms regulating tissue-specific chromosome arm-wide gene expression include DNA methylation, histone acetylation and higher order nuclear organization, the 4D Nucleome^23^. Data for chromosome-specific histone acetylation and nuclear organization are not readily available, but the Gene Expression Omnibus (GEO) database provides genome-wide methylation patterns. Based on these data we analyzed chromosome arm-wide methylation patterns for 11 tissues from 765 samples (Materials and Methods). While differences in the methylation status of specific genes and their expression were readily apparent, arm-level differences in methylation levels did not allow the discernment of specific tissues from each other (Supplementary Fig. 3) and could not explain the chromosome-arm wide tissue-specific gene expression patterns.

## DISCUSSION

Overall, our results indicate that differences of chromosome arm-wide gene expression levels in normal human tissues are enhanced by the acquisition of aneuploidies in the cognate tumors, suggesting a non-oncogenic, tissue-specific physiological basis for clonal expansion. Interestingly, Sack and colleagues^24^ have demonstrated that the inclusion of tissue-specific growth promoting genes strengthens the correlation between chromosome arm loss/gain ratios and the proliferation-driving capability of each chromosome-arm in breast and pancreatic cancers. A general, yet not tissue-specific, role of copy number alterations and metabolic selection pressure was reported by Graham and colleagues^25^. Of note, several publications point to a reduction of cellular fitness as a consequence of general aneuploidy^26–28^. We show that, unlike general aneuploidy, tissue-specific aneuploidies that enhance chromosome arm-wide normal tissue-specific gene expression levels result in clonal expansion. Notably, we previously showed that the gain of chromosome 13 in colorectal cancer activates both Notch and Wnt signaling^29^, and that the acquisition of extra copies of chromosome 7 results in upregulation of the Wnt pathway (Braun et al., accepted for publication, Neoplasia), which supports our finding that the enhancement of tissue-type specific chromosome arm-wide gene expression levels by copy number alterations can promote cellular fitness.

In conclusion, we found that (i) chromosome arm-wide gene expression patterns are tissue-type specific and predict the respective normal tissue of origin. (ii) The acquisition of cancer-type specific aneuploidies mirrors chromosome arm-wide gene expression patterns of the respective *normal* tissues, i.e., the changes are inherited after cell division and become “hardwired”. Our observations are schematically summarized in Fig. 4. Rather than being acquired and maintained based on an *a priori* oncogenic advantage, the dominant and ubiquitous cancer-type specific chromosomal aneuploidies are under strong selection to optimally fit the physiological transcriptional requirements of their respective normal tissues.

**Figure 4:**
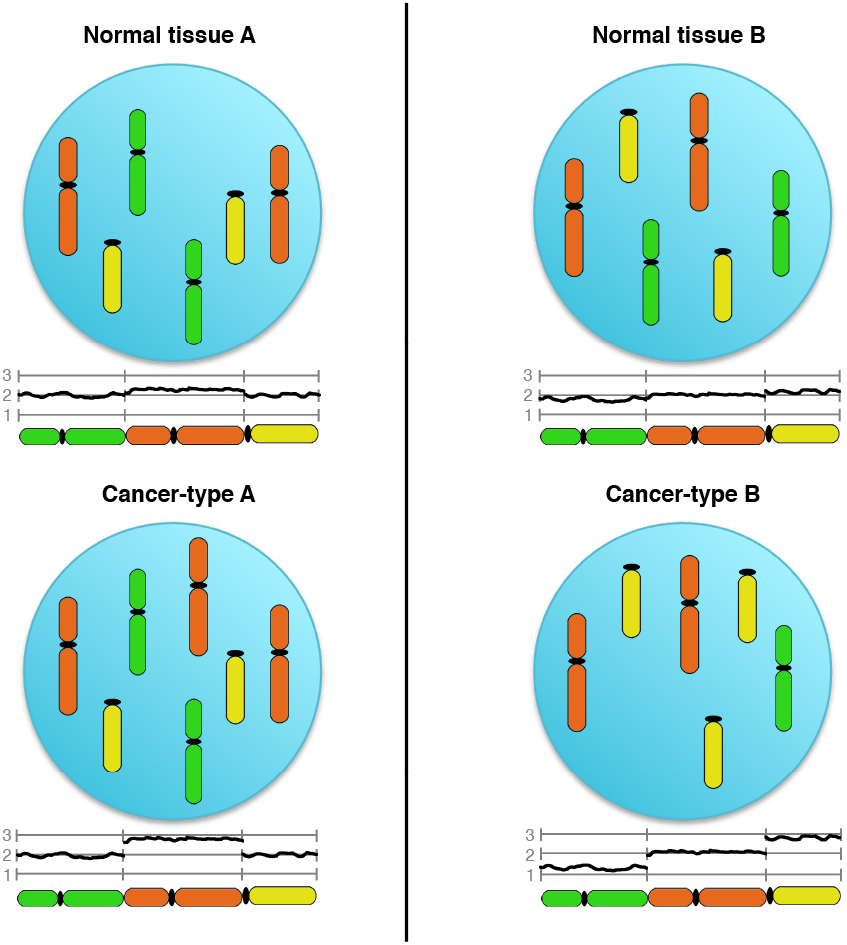
Schematic presentation of the results. Genes on the red chromosomes are expressed at slightly higher levels compared to other chromosomes in normal tissue A, whereas in normal tissue B, the yellow chromosomes shows increased tissue-specific expression and genes on the green chromosome are expressed at lower levels. This results in a subtle increase or decrease in chromosome arm-wide transcript levels, respectively. The acquisition of chromosomal aneuploidies in the respective tumors (gain of the red chromosome in tumor A and the yellow chromosome in tumor B, accompanied by the loss of the green chromosome in tumor B) amplifies this effect and provides the genetic basis of “hard-wiring” tissue-specific chromosome arm-wide gene expression levels as the basis for clonal expansion.

## MATERIALS AND METHODS

### Tissue and tumor type inclusion

In this study, the cancer types and respective normal tissues included were all those for which there was availability of (1) genomic copy number of the cancerous tissue from The Cancer Genome Atlas (TCGA) and (2) gene expression of normal tissues from Genotype-tissue Expression (GTEx) Database. Epithelial tumor types with no chromosomal aneuploidy (i.e. no chromosomal arm was gained or lost in 25% of the samples, as defined throughout this study, see below) were excluded from the analysis. For tissues with more than one matching cancer type in TCGA (e.g., lung), the cancer type with the larger number of samples with somatic copy number alteration data was selected (e.g., TCGA LUAD rather then LUSC).

### Computation of chromosomal arm gain and loss score in cancerous tissues

We used the TCGA sample-wise chromosomal arm gain and loss data provided^12^, where the ploidy was determined via the ABSOLUTE algorithm^30^. Each segment was designated as amplified, deleted, or neutral compared to the ploidyof each sample. Tumors altered <20% were considered “non-aneuploid,” and others were designated “other.” The scores of each arm are −1 if lost, +1 if gained, 0 if non-aneuploid, and “NA” otherwise.

For each of the 39 chromosomal arms we define an arm aneuploidy score for each cancer type by subtracting the number of chromosomal arm losses from the number of chromosomal arm gains and normalized by the sample size. Formally:

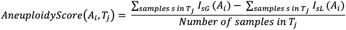

Where *A_i_* is chromosomal arm *i* (of 1-39 chromosomal arms), *T_j_* is tumor type *j* (of overall 15 tumor types considered) and the indicators *I_sG_* (*A_i_*) and *I_sL_* (*A_i_*) are defined by:

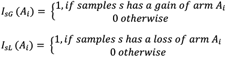

### Computation of chromosome-arm level gene expression changes in normal tissues

We count the number of up and down regulated genes in each arm, for every cancer type, using the following:

1. Up regulated genes: for each chromosomal arm, we first identify the cancer types that are gaining this arm (if more than 25% of the TCGA samples of that cancer type has a gain of the arm) and those that do not (all other cancer types). Then, for each normal tissue corresponding to a cancer type with a gain of that arm, we find the genes (on that chromosomal arm) which are up-regulated vs. the normal tissues corresponding to tumors with no gain of that arm, using one-sided Rank-sum test P-value<0.05. Similarly, for each normal tissue corresponding to a cancer type with no gain of the arm, we find the genes (on that chromosomal arm) which are up-regulated vs. normal tissues corresponding to tumors with gain of that arm, using one-sided Rank-sum test P-value<0.05.
2. Down regulated genes: similarly, for each chromosomal arm, we identify the cancer types that are losing that arm (if more than 25% of the TCGA samples of that cancer type has a loss of the arm) vs. those that do not. For each normal tissue corresponding to a cancer type with a loss of the arm, we then find the genes (on that arm) which are down-regulated vs. normal tissues corresponding to tumors with no loss of that arm, using onesided Rank-sum test P-value<0.05. Similarly, for each normal tissue corresponding to a cancer type with no loss of the arm, we find the genes (on that arm) which are down-regulated vs. normal tissues corresponding to tumors with loss of that arm, using onesided Rank-sum test P-value<0.05.

Then, the arm-level gene expression regulation score is defined by subtracting the number of down-regulated genes from the number of up-regulated genes on each chromosomal arm, for each tumor type, which is then normalized by the number of genes lying on each arm. Formally:

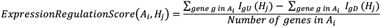

Where *A_t_* is chromosomal arm *i* (of 1-39 chromosomal arms), *H_j_* is normal tissue *j* (of overall 15 normal tissues considered) and the indicators *I_gU_* (*H_j_*) and *I_gD_* (*H_j_*) are defined by:

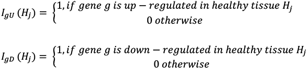

### Correlation of arm level regulation across normal tissues

We evaluate the correlation between the arm gain/loss score in each cancer type and the score of up/down-regulation in each normal tissues via two approaches (for both using Spearman rank correlation rho and P-value):

1. Arm-level correlation: for each chromosomal arm, we correlate the number of samples with gain of that arm for each cancer type, with the number of up regulated genes in that arm for the corresponding normal tissues. Similarly, for each arm we correlate the number of samples with loss of that arm for each cancer type, with the number of down regulated genes in that arm of the corresponding normal tissues.
2. Tumor/normal tissue correlation: For each tumor type considered, we correlate the arm aneuploidy score with the arm gene expression regulation score of the respective normal tissue.

### Normal tissue classification

To classify normal tissues using the chromosomal-arm regulation map of those tissues, we calculate for each sample, the median gene expression level of the genes in each chromosomal arm. We then perform K-Nearest-Neigbors (KNN, with K=5) classification with a Leave-One-Out cross validation (LONCOV), aiming to classify each sample by the median chromosomal arm expression of the normal tissues that are closest to it, and calculate the resulting accuracy (percentage of correctly classified samples in the LONCOV). For comparison, we perform a similar KNN analysis with the full gene expression data.

### Evaluating the correlation between arm-level gain and loss in tumors with the localization of oncogenes and tumor suppressor genes

We downloaded oncogene and tumor suppressor classifications for each gene in each tissue from^31^. Then, for each arm and tissue, we counted the number of genes on that arm classified as oncogene (respectively tumor suppressor) resulting in a tissue specific oncogene (respectively, tumor suppressor) enrichment profile for the arm. Finally, to evaluate whether an arm is gained more often in tissues where it harbors oncogenes (respectively, lost more often in tissues where it harbors tumor suppressors) we measure the spearman rank correlation between the tissue-specific gain (respectively, loss) frequency of that arm and its tissue specific oncogene (respectively, tumor suppressor) enrichment profile.

### Tissue specific methylation analysis

We curated a list of 18 Illumina 450K methylation datasets from Gene Expression Omnibus covering tissues from 11 organs. These datasets span different studies comparing methylation profiles of tissues between diseased and normal control individuals. We only selected methylation profiles of normal control individuals for further analysis. Multiple datasets containing samples coming from the same tissue were merged to generate one methylation dataset. (See Supplementary Table 1 for more information on each of the datasets). In order to do a comparison of methylation levels on each arm across tissues, we first pre-process each dataset in the following three steps.

1. Filtering out probes overlapping with single nucleotide polymorphisms to control for population specific differences in methylation levels^32^.
2. Rescaling the beta values between type 1 and type 2 probes using beta mixture quantile normalization. This minimizes technical differences that may arise between two different probe designs^33^.
3. Rescaling beta values of each sample in a dataset by the median value of the dataset to adjust for dataset-specific differences in methylation levels.

Given a chromosome arm, we find each tissue where that arm is gained in the corresponding cancer (as defined above). We measure the fold change in methylation levels of each probe on that arm in that tissue, relative to tissues where that arm is not gained. Likewise, we find each tissue where the arm is lost in the corresponding cancer (as defined above). We measure the fold change in methylation levels of each probe on that arm in that tissue, relative to tissues where that arm is not lost. Finally, we find each tissue where the arm is neither gained nor lost in the corresponding cancer. We measure the fold change in methylation levels of each probe on that arm in that tissue, relative to tissues where the arm is either gained or lost.

We define *I_pU_* (*H_j_*) = 1, if fold change of probe *p* located on chromosome arm *A_i_* in tissue *H_j_* > 2 (hypermethylation). Similarly, *I_pD_* (*H_j_*) = 1 if the fold change of probe *p* in tissue *H_j_* < 1/2 (hypo-methylation). The tissue-specific methylation score of arm *A_i_* in tissue *H_j_* is then evaluated as follows:

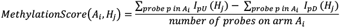

To assess whether chromosome arm level differences in gene expression of different normal tissues can be explained by chromosome arm level differences in methylation, we measure the spearman rank correlation between number of downregulated genes on an arm in a tissue and number of hypermethylated probes (rho-). This follows from the observation that genes in close genomic proximity to hypermethylated regions are less likely to be expressed. Likewise, we also measure the spearman rank correlation between number of upregulated genes on an arm in a tissue (see above) and number of hypomethylated probes (rho+). The results of this analysis are shown in Supplementary Figure 3.

### Inferring which arm aneuploidies occur early in tumorigenesis

For each of the 15 tissues considered, given the aneuploidy profiles of 39 chromosome arms in corresponding primary tumors of different patients, one can partially order these aneuploidies by time using the following score:

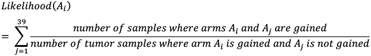

This score estimates the likelihood of occurrence of an event given the occurrence of other events. It has been previously shown that the lower this score, the earlier the event^34^. Given that tumors of different tissues may evolve at different rates, we defined an event as *early* if the event was among the *first 10* events after sorting them in increasing order by the above score. Our downstream analyses are robust at more stringent definitions of an early event (i.e., first 9, first 8, first 7, …, first 2, first event; results not shown).

## ACKNOWLEDGEMENTS

The authors are indebted to Drs. Thomas Cremer, Marion Cremer, Reinhard Ebner, Kenneth C. Carter, W. Michael Kuehl, Javed Khan, Alejandro Schäffer and E. Michael Gertz for valuable comments on the manuscript and to Buddy Chen for editorial assistance. The study was supported by the Intramural Research Program, National Cancer Institute/NIH. DH and RB were supported by the Deutsche Krebshilfe, GE through the Deutsche Forschungsgemeinschaft, NA through the NCI/University of Maryland Graduate Partnership Program, and DB by a Wellcome Trust/NIH PhD Studentship. The results published here are in part based upon data generated by the TCGA Research Network: http://cancergenome.nih.gov/.

---

**Figure S1:**
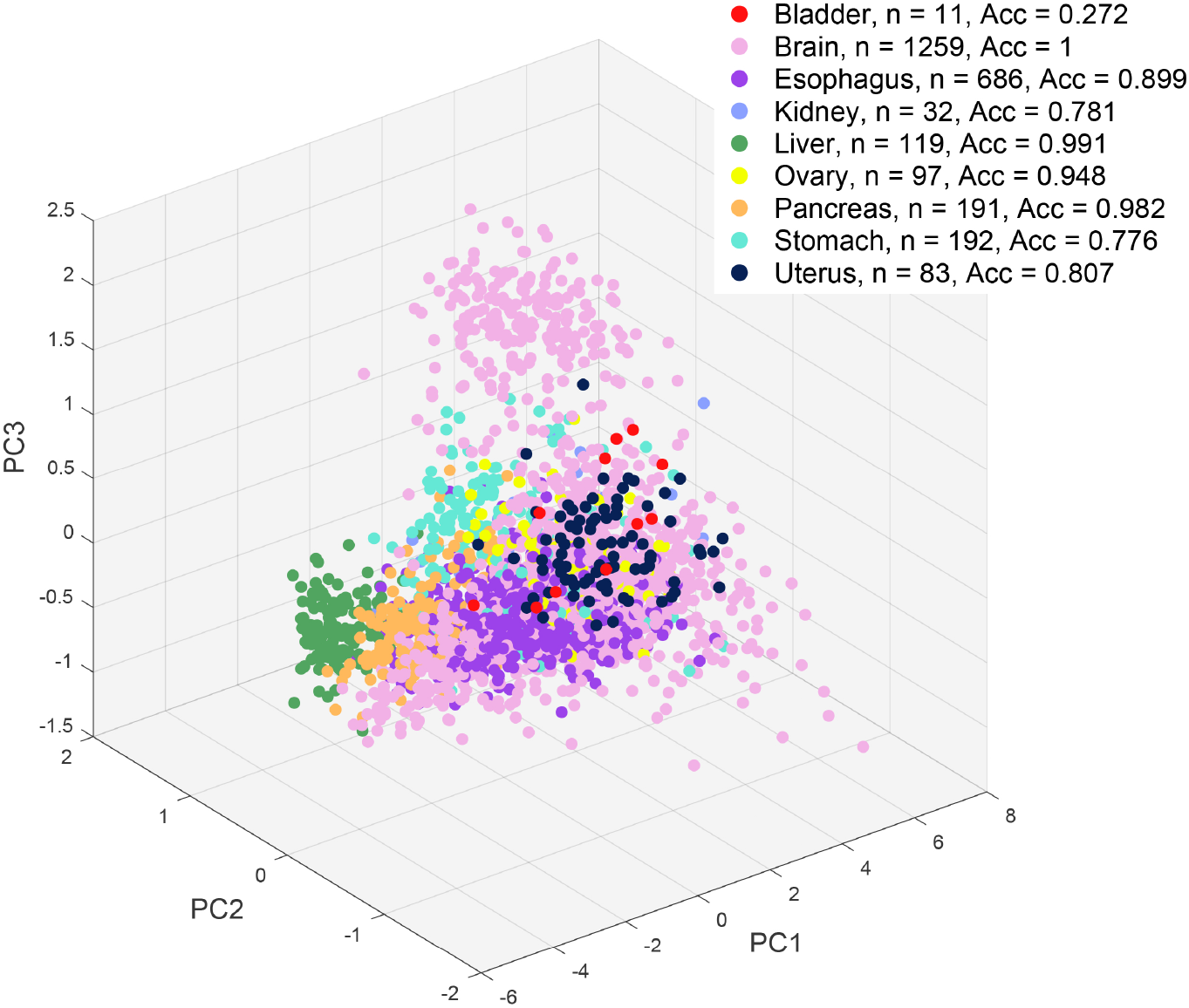
Chromosome arm-wide gene expression profiles allow tissue classification. The PCA analysis of chromosome arm-wide gene expression of normal tissues is displayed for nine additional tissues analyzed. The different normal tissues can be predicted with high accuracy.

**Figure S2:**
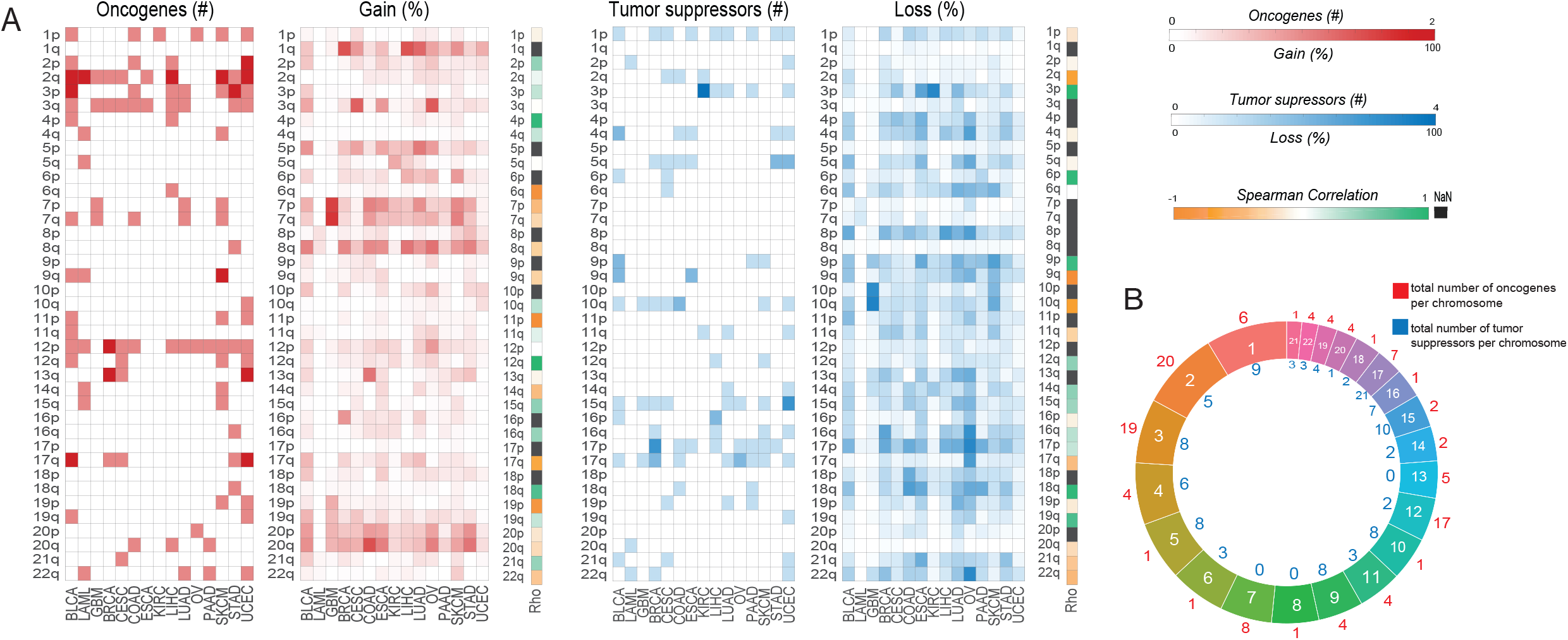
(A) Correlation of oncogenes and tumor suppressor genes with the gains and losses of chromosome arms, respectively, on which they reside. The correlation does not reach significance. BLCA, bladder cancer; LAML, acute myelocytic leukemia; GBM, glioblastoma multiforme; BRCA, breast cancer; CESC, cervical squamous carcinoma; COAD, colorectal adenocarcinoma; ESCA, esophageal squamous carcinoma; KIRC, renal cell carcinoma; LIHC, hepatocellular carcinoma; OV, ovary carcinoma; PAAD, pancreatic adenocarcinoma; SKCM, melanoma; STAD, stomach adenocarcinoma; UCEC, uterine endometrial carcinoma. (B) The circle indicates the distribution of the number of oncogenes and tumor suppressor genes on each chromosome and is the basis for the results presented in the Figure.

**Figure S3:**
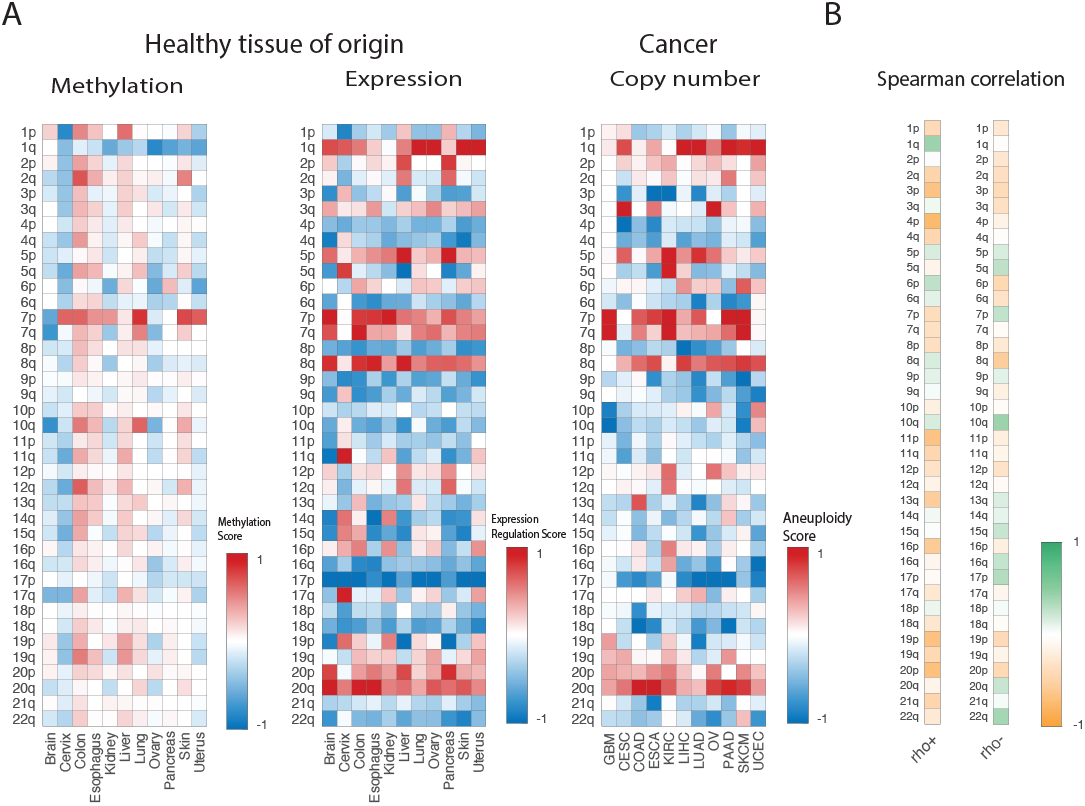
(A) Chromosome arm-wide methylation, chromosome arm-wide gene expression in normal tissues, and cancer type-specific chromosomal aneuploidy patterns in the respective tumors. (B) The correlations of the arm-wide methylation and gene expression in normal tissues are not significant.

**Supplementary Table 1:** Methylation Datasets

| **Supplementary Table 1:**<br>**Methylation Datasets** |  |  |  |
| --- | --- | --- | --- |
| **Dataset ID** | **Platform** | **Tissue** | **Brief Description** |
| GSE32146 | Illumina 450K | Colon | 10 normal Colon mucosa tissue, 10 Crohn's disease, 4 ulcerative colitis |
| GSE40360 | Illumina 450K | Brain, frontal lobe | 47 samples analyzed in total: 17 MS females, age $55.5 \pm 10.3$ years; 11 MS males, age $55.0 \pm 9.9$ years; 7 control females, $69.1 \pm 9.2$ years; 12 control males, $64.7 \pm 8.0$ years |
| GSE61107 | Illumina 450K | Brain, frontal cortex | Genome-Wide DNA methylation analysis was performed on post-mortem human brain tissue from 24 patients with schizophrenia and 24 unaffected controls |
| GSE88890 | Illumina 450K | Brain, cortex | Tissue (n=75) from two regions of the cortex (Brodmann area 11 (BA11, n=40) and Brodmann area 25 (BA25, n=35)) from 20 MDD MDD suicide cases and 20 non-psychiatric sudden death controls |
| GSE89702 | Illumina 450K | Brain, cerebellum | 33 post-mortem brain (cerebellum) samples were obtained from the Douglas Bell-Canada Brain Bank (DBCBB; 16 schizophrenia and 17 controls), Montreal, Canada |
| GSE89703 | Illumina 450K | Brain, hippocampus | 27 post-mortem brain (hippocampus) samples were obtained from the London Brain Bank for Neurodegenerative Disorders (LBBND; 14 schizophrenia and 13 controls), London, UK |
| GSE89705 | Illumina 450K | Brain, Striatum | 33 post-mortem brain (striatum, putamen) samples were obtained from the Douglas Bell-Canada Brain Bank (DBCBB; 16 schizophrenia and 17 controls), Montreal, Canada |
| GSE62640 | Illumina 450K | Pancreas | 53 male and 34 female human pancreatic islet samples. Normally methylated, non-methylated and fully methylated human DNA samples were included as controls |
| GSE51954 | Illumina 450K | Skin | 10 younger sun protected dermal samples, 10 younger sun exposed dermal samples, 10 older sun protected dermal samples, 10 older sun exposed dermal samples, 9 younger sun protected epidermal samples, 9 younger sun exposed epidermal samples, 10 older sun protected epidermal sample, 10 older sun exposed epidermal samples |
| GSE90124 | Illumina 450K | Skin | Samples: Bisulphite converted DNA from the 322 samples were hybridised to the Illumina HumanMethylation450 BeadChip |
| GSE52401 | Illumina 450K | Lung | Bisulphite converted DNA from 244 fresh-frozen primary human lung samples were hybridised to the Illumina Infinium 450k Human Methylation Beadchip v1.2. This file has 60 samples. |
| GSE61258 | Illumina 450K | Liver | Bisulphite converted DNA from the 79 samples were hybridised to the Illumina Infinium 450k Human Methylation Beadchip |
| GSE61446 | Illumina 450K | Liver | Bisulphite converted DNA from the 67 Liver samples were hybridised to the Illumina Infinium 450k Human Methylation Beadchip |
| GSE51820 | Illumina 450K | Ovary | This study included 96 total samples, including 53 serous ovarian cancers, 11 endometrioid ovarian cancers, 13 clear cell ovarian cancers, 8 mucinous ovarian cancers, 4 fallopian tube epithelium specimens, 4 ovarian surface epithelium specimens, 1 universally methylated DNA sample, 1 normal human lymphocyte sample, and 1 50:50 mixture of the normal human lymphocyte DNA and universally methylated DNA samples |
| GSE46306 | Illumina 450K | Cervix | In this study 20 normal cervical samples (HPV negative), 18 samples with CIN3 lesions (HPV positive) and 6 cervical cancer tissues (HPV positive) were included. |
| GSE45187 | Illumina 450K | Uterus | Bisulphite converted DNA from the three uterine leiomyoma, three myometrium with leiomyoma and three myometrium without leiomyoma were hybridised to the Illumina Infinium HumanMethylation450 BeadChip. |
| GSE52826 | Illumina 450K | Esophagus | DNA methylation profiles of esophageal squamous cell carcinoma (4 samples), paired adjacent normal surrounding tissues (4 samples) and normal esophagus mucosa from healthy individuals (4 samples) were generated using Infinium methylation 450K BeadChips from Illumina (Illumina, San Diego, USA) |
| GSE59157 | Illumina 450K | Kidney | Genome wide DNA methylation profiling of normal kidney (n=36), nephrogenic rest (n=22) and Wilms tumour (n=37) was performed using the Illumina 450k array. |

